## Supplementary materials and methods for "Upregulated expression of ubiquitin ligase TRIM21 promotes PKM2 nuclear translocation and astrocyte activation in experimental autoimmune encephalomyelitis"

**Supplementary Table S1. Primers used in this study.**

| Name | Primer sequences (5'-3' orientation) |
|---|---|
| PKM2 | Forward: GCCGCCTGGACATTGACTC |
| | Reverse: CCATGAGAGAAATTCAGCCGAG |
| TRIM21 | Forward: GGGAGGAGGTCACCTGTTCTA |
| | Reverse: GGCACTCGGGACATGAACTG |
| IL-6 | Forward: GCTGGAGTCACAGAAGGAGTGGC |
| | Reverse: GGCATAACGCACTAGGTTTGCCG |
| IL-1β | Forward: CACTACAGGCTCCGAGATGAACAAC |
| | Reverse: TGTCGTTGCTTGGTTCTCCTTGTAC |
| TNF-α | Forward: CCTGTAGCCCACGTCGTAG |
| | Reverse: GGGAGTAGACAAGGTACAACCC |
| Cyclin D1 | Forward: AAGTGCGTGCAGAAGGAGATTGT |
| | Reverse: GGATAGAGTTGTCAGTGTAGATGC |
| GAPDH | Forward: AGGTCGGTGTGAACGGATTTG |
| | Reverse: TGTAGACCATGTAGTTGAGGTCA |

### Supplementary Figures

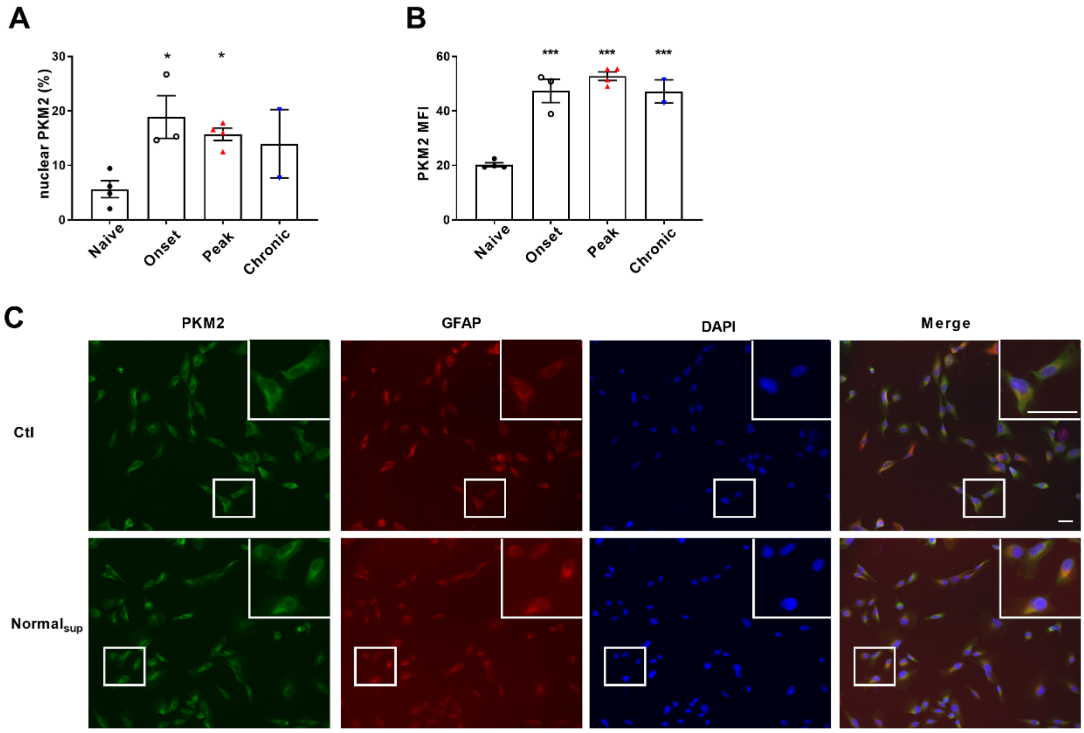


**Figure S1. Quantification of nuclear ratio of PKM2 in astrocytes and mean fluorescence of PKM2 in control and EAE mice.** (A) Nuclear PKM2 ratio in control mice and different phases of EAE mice were calculated in Figure 1A. Number of nuclear PKM2 in astrocytes was quantified by Imgae-Pro plus manually (eg. PKM2 was only counted in GFAP$^+$ astrocytes cells, nuclear or cytoplasmic based on DAPI blue staining). The proportion of nuclear PKM2 in astrocytes is determined by normalizing the count of nuclear PKM2 to the count of GFAP$^+$ cell numbers. (B) Mean fluorescence intensity (MFI) of PKM2 in control and EAE mice was calculated by Image J. (C) Immunofluorescence staining of PKM2 (green) with GFAP (red) in non-treated primary astrocytes (control) or primary astrocytes cultured with splenocytes supernatants from normal mice (Normal$_{sup}$). Scale bar: 50 µm. Data are represented as mean ± SEM, one-way ANOVA, **P*<0.05; ****P*<0.001. SEM, standard error of the mean.

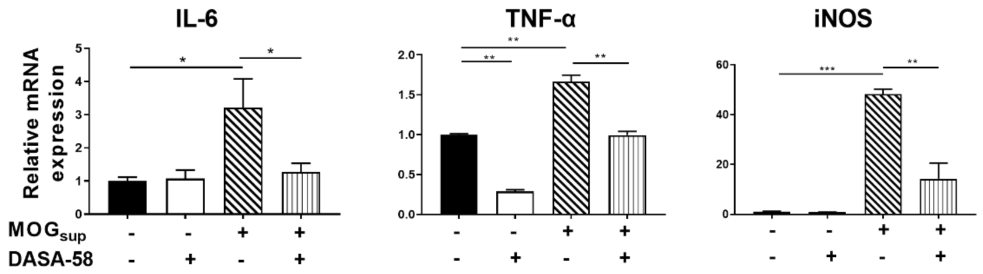


**Figure S2. qPCR analysis of mRNA levels of inflammatory cytokines.** Primary astrocytes were pretreated with 50 μM DASA-58 for 30 min and stimulated with $MOG_{sup}$ for 12h. Data are represented as mean ± SEM, one-way ANOVA, **P*<0.05; ***P*<0.01; ****P*<0.001. SEM, standard error of the mean.

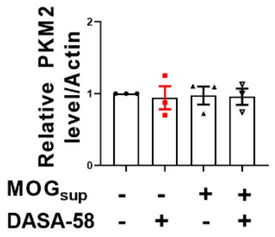


**Figure S3. Quantification of PKM2 protein level in astrocytes treated with $MOG_{sup}$ or $MOG_{sup}$ pretreated with DASA-58.** Protein levels were calculated by Image J, PKM2 expression was normalized to β-Actin level. One-way ANOVA, data are represented as mean ± SEM.

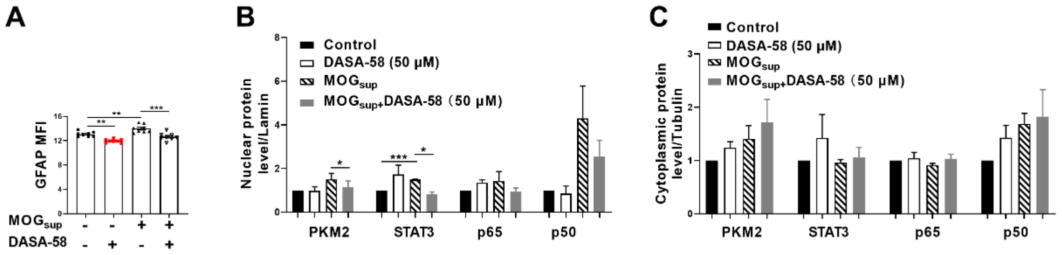


**Figure S4. Quantification results of GFAP and cyto-nuclear protein levels in astrocytes treated with $MOG_{sup}$ or $MOG_{sup}$ pretreated with DASA-58.** (A) MFI analysis of GFAP in different groups. Primary astrocytes were pretreated with 50 μM DASA-58 for 30 min and stimulated with $MOG_{sup}$ for 12h. (B) Quantification of nuclear protein levels of PKM2, p50, p65 and STAT3 in different groups. The nuclear levels of the indicated proteins were normalized to lamin. (C) Quantification of cytoplasmic protein levels of PKM2, p50, p65 and STAT3 in different groups. The cytoplasmic levels of the indicated proteins were normalized to tubulin. Data are represented as mean ± SEM, one-way ANOVA, **P*<0.05; ***P*<0.01; ****P*<0.001.

**A**

| Gene | Protein names | Unique peptides (Ctl) | Unique peptides ($MOG_{sup}$) |
|---|---|---|---|
| Pkm | Pyruvate kinase PKM | 4 | 5 |
| Anxa2 | Annexin A2 | 4 | 4 |
| Eno1 | Alpha-enolase | 4 | 1 |
| Ckmt1 | Creatine kinase U-type, mitochondrial | 3 | 0 |
| Ckb | Creatine kinase B-type | 2 | 1 |
| Aldoa | Fructose-bisphosphate aldolase A | 2 | 3 |
| Mdh2 | Malate dehydrogenase, mitochondrial | 1 | 2 |
| Slc25a4 | ADP/ATP translocase 1 | 2 | 2 |
| Ldha | L-lactate dehydrogenase A chain | 2 | 0 |
| Ldhc | L-lactate dehydrogenase C chain | 2 | 0 |
| Fabp5 | Fatty acid-binding protein 5 | 1 | 1 |
| Trim21 | E3 ubiquitin-protein ligase TRIM21 | 0 | 1 |

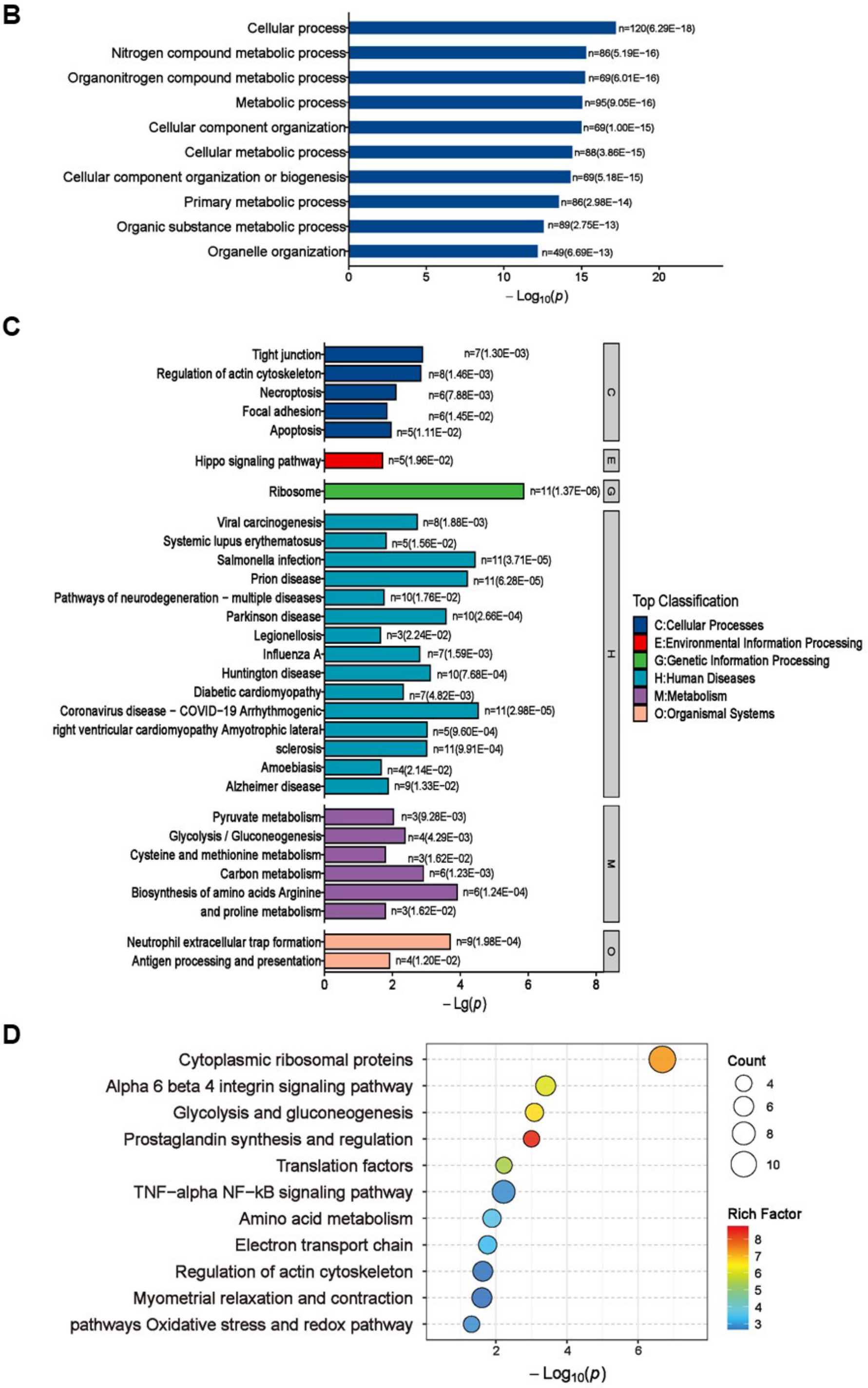

**Figure S5. Mass spectrometry results of PKM2-interacting proteins in astrocytes.** (A) Mass spectrometry (MS) showed the list of metabolic-related proteins that potentially interact with PKM2 in unstimulated (Ctl) and $MOG_{sup}$-stimulated primary astrocytes. TRIM21 was identified to interact with PKM2. (B-D) Biological process of GO term (B), KEGG pathway (C) and Wikipathway (D) analysis of proteins identified by MS.

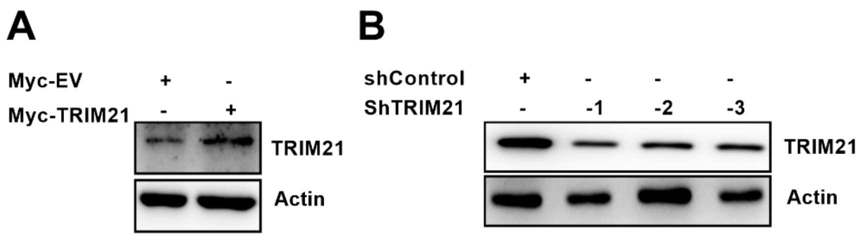


**Figure S6. Verification of TRIM21 overexpression and knockdown efficiency.** (A) Overexpression of TRIM21 was verified by western blotting analysis. (B) Western Blotting analysis of TRIM21 knockdown efficiency. Sh: short hairpin; EV: empty vector.

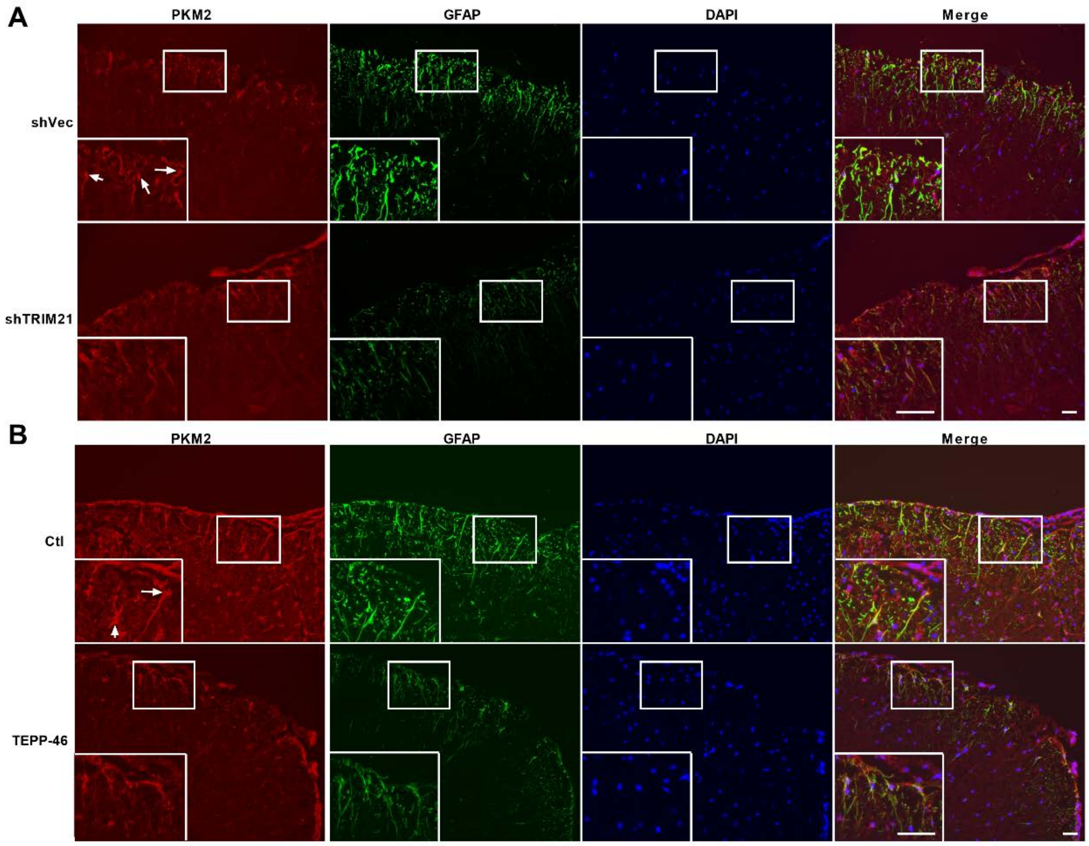


**Figure S7. PKM2 expression and localization in shTRIM21-treated and TEPP-treated EAE mice.** (A) PKM2 expression in spinal cord of mice from shVec and shTRIM21-treated EAE mice was measured by immunofluorescence. (B) PKM2 expression in spinal cord of mice from TEPP-46- or vehicle-treated EAE mice was measured by immunofluorescence. White arrows indicate nuclear PKM2. DAPI (blue) was used as a nuclear staining. Scale bar: 50 μm.

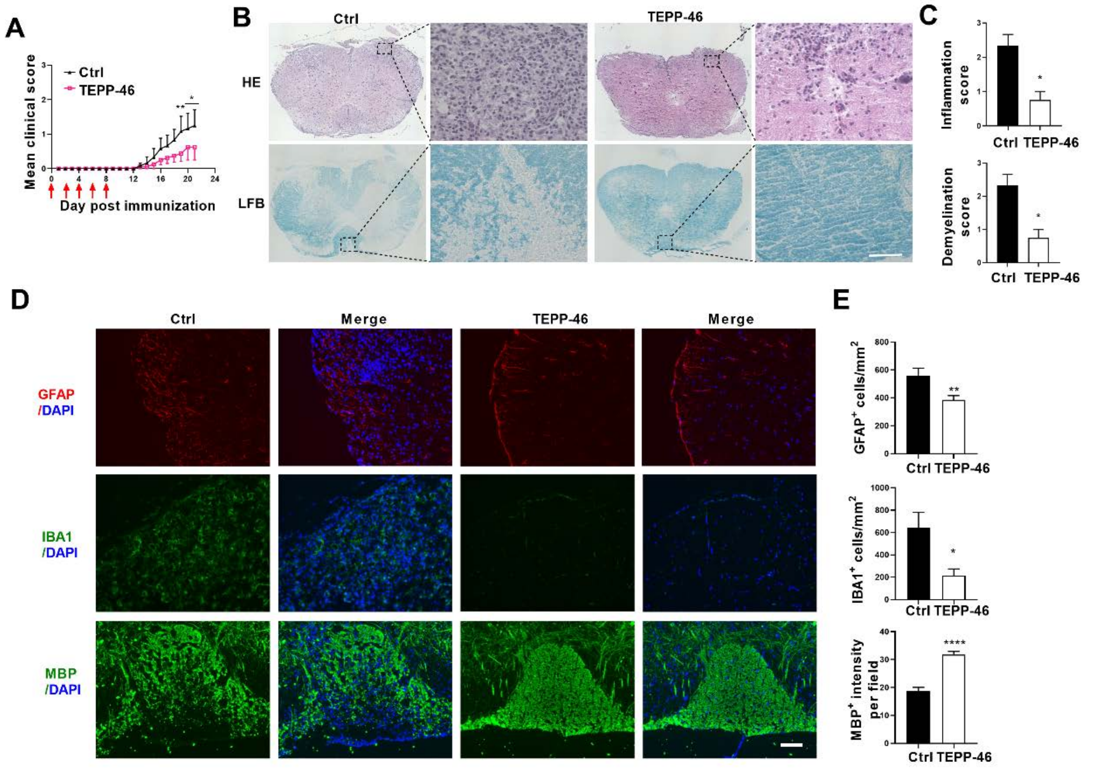


**Figure S8. i.p. injection of TEPP-46 alleviated the development of Experimental Autoimmune Encephalomyelitis (EAE).** C57BL/6 mice were i.p injected with 200 μl vehicle or 50 mg/kg TEPP-46 dissolved in vehicle every other day from day 0 to day 8 p.i.. Mice were sacrificed at day 21 p.i. and spinal cords were harvested. (A) Disease was scored daily on a 0 to 5 scale. N=6 to 8 mice in each group. (B) Spinal cord sections were stained for markers of inflammation by hematoxylin and eosin (H&E) and demyelination by Luxol fast blue (LFB), respectively. Scale bar: 50 μm. (C) Scoring of inflammation (H&E) and demyelination (LFB) on a 0-3 scale. (D) Immunostaining of GFAP, IBA1 and MBP on spinal cord sections of TEPP-46- or vehicle-treated EAE mice. (E) Quantification of GFAP positive cells/mm$^2$, IBA1 positive cells/mm$^2$ in the white matter of the spinal cord. MBP intensity was measured in the white matter of the spinal cord using Image-Pro. The measured areas included 3 to 5 fields per group. i.p., intraperitoneally; p.i., postimmunization; Scale bar: 100 μm. Data are represented as mean ± SEM. **P*<0.05; ***P*<0.01; ****P*<0.001, as determined by two-way ANOVA analysis (A) or unpaired Student's t test (C, E).
